# Identification and Assessment of Systematic Measurement Error on Electrophysiological Recordings of Neural Cell Cultures

**DOI:** 10.1101/2022.05.27.493606

**Authors:** Tyler Stone, Thomas R. Kiehl, Charles Bergeron

## Abstract

Microelectrode arrays (MEA) hold great promise for a broad range of applications that require reliable characterization of the growth and function of neurons in culture. Widespread adoption of this platform depends on analytical methods to extract meaning from highly variable and noisy observations. In analyzing a comprehensive database of MEA recordings, we discovered that 22% of the electrodes presented systematic patterns of under- or non-detection of spike activity. Going undetected, principal components analysis (PCA) of these data reveal trends that would have lead to incorrect biological interpretations. We fully document these *defective* or biased electrodes, and distinguish two forms of defectiveness, via representations that aid in detecting them. We also showcase our approach for analyzing these data that permit for post-analytic review and correction. Repeating our PCA on *cleaned* data, we discover a more complex interplay of biological variability. Finally, we make a case for transparency in data reporting and propose best practices for experimental and analysis phases.

## I. Introduction

**M**ULTIELECTRODE arrays (MEA) are broadly used for in vitro neural cell culture observation [1]–[7]. An MEA consists of a glass plate with electrodes that record action potentials (voltage dynamics) from nearby cells, typically neurons. The recorded voltages are then fitted to locate stereotypical *spikes* indicating the activity of the observed cell population. Spike patterns are then analyzed, under the hypothesis that valuable biological insights can be derived from these data.

Recently, several commercial MEA platforms have come to market as researchers seek to further quantify the behavior of their neural cell cultures. Complete realization of these platforms’ potential to the study of disease and development depends on the capability to

- make reliable and repeatable observations,
- characterize neuronal network behavior, and
- make quantitative comparisons between cultures.

These observations, characterizations and comparisons must be independent of the inherent randomness of network structure and variation in recording procedures. In this paper, we demonstrate that naïve processing of these data leaves us vulnerable to bias.

Previously, Waagenar et al. [8] published an extensive set of recordings performed on cultures growing on 60-electrode MEA’s. Their MEA platform includes an off-the-shelf amplifier and open-access software [9], [10]. Their goal was to learn about the dissociated embryonic rat cortex cells that they cultured, classifying each recording in terms of burst shape and frequency, finding a strong influence in between plating density and spike activity, and characterizing the baseline spike activity with developmental age [8].

To promote the value of this platform, they made their 86-gigabyte spike train dataset available to the research community. This paper re-examines these data. Surprisingly, few researchers have analyzed these data: there have been a few follow-up papers from their group [11]–[13], a few applications of network modeling to selected recordings [14], [15], and a detailed study of neuronal connectivity evolution based on the false discovery rate [16]. We identify two sources of systematic measurement error that affected 13 electrodes. This bias may have confounded previous analysis attempts. We highlight possibilities for post-experimental review to identify and handle these issues. We also make a case for transparency in data reporting and propose best practices for experimental and analysis phases.

By design, each electrode simultaneously records its electrical activity from a small proximal subpopulation of neural cells within a culture. While MEA’s don’t reach the same specificity or fidelity of single-cell techniques, such as patch-clamp recordings, they do allow for multiple simultaneous recordings from a cell population. Yet, recordings are usually assessed by counting spikes detected across all electrodes per unit time: the array-wide spike detection rate (ASDR) [8]. In doing so, the contributions of all electrodes are merged into a single feature. This state-of-the-art has persisted for over a decade. In this paper, we show that we can glean additional insights by treating each electrode’s spike detection rate (SDR) as a distinct feature. Then, we handle the increase in the number of features by performing principal components analysis (PCA) to identify dominant trends across the electrodes. We determine that ASDR captures approximately 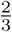 of the overall variability, and we present ways of visualizing dominant trends in the remaining variance. In doing so, we identify previously-unknown recording artifacts on an established, large, public dataset, and establish that meaningful information is lost when all electrodes are aggregated. Thereupon, we contribute towards making MEA recordings of electrophysiological activity an increasingly meaningful asset towards studying neuron development in culture.

## II. Notation

Let *X* denote a matrix of real numbers, and let *X*_*i,j*_ denote the entry of *X* appearing in its *i*th row and *j*th column. The transpose of *X* is *X*^*T*^. Let 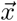 denote a vector, and let 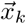 be the *k*th column of matrix *X*. The scalar product of 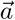 and 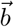 is written 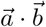 while the Euclidean norm of 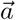 is 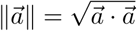.

The singular value decomposition (SVD) of an *ℓ*-by-*n* matrix *X* is

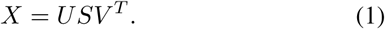

In this paper, *n* < *ℓ* so that

- *U* is an *ℓ*-by-*n* orthonormal matrix whose columns 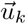 are the left singular vectors of *X*,
- *S* is an *n*-by-*n* diagonal matrix whose non-negative entries *S*_*k,k*_ are the singular values of *X* sorted in decreasing order, and
- *V* is an *n*-by-*n* orthonormal matrix whose columns 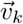 are the right singular vectors of *X* [17].

In some contexts, we explicitly state that matrix *X* possesses *n* columns using a square-bracketed superscript: *X*^[*n*]^. It’s SVD is then expressed as

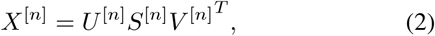

where, *U* ^[*n*]^ is the matrix of left singular vectors decomposed from the *n*-column matrix *X*^[*n*]^, and 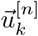 is the *k*th left singular vector of *X*^[*n*]^. Later in this paper, this heavy notatation will be particularly useful when comparing decompositions (equation 7).

## III. Methods

We summarize the previously-described dataset of Wagenaar et al. [8]. They used MEAs (Multichannel Systems, Reutlingen, Germany) consisting of *n* = 59 transparent electrodes arranged in an 8-by-8 square grid with the four corners missing and one electrode on the array’s edge replaced by a large ground electrode. Figure 1 helps us visualize this configuration. The electrode diameter is 30 *µ*m. Electrodes arespaced 200 *µ*m apart. A *batch* consists of embryonic cortex cells (neurons and glia) dissected from the litter of a pregnant Wistar rat at day 18 of gestation. Cells were dissociated by sequential application of papain and trituration, and then plated on MEAs so as to observe the re-emergence of neuronal interconnectivity in these cultures.

**Fig. 1.**
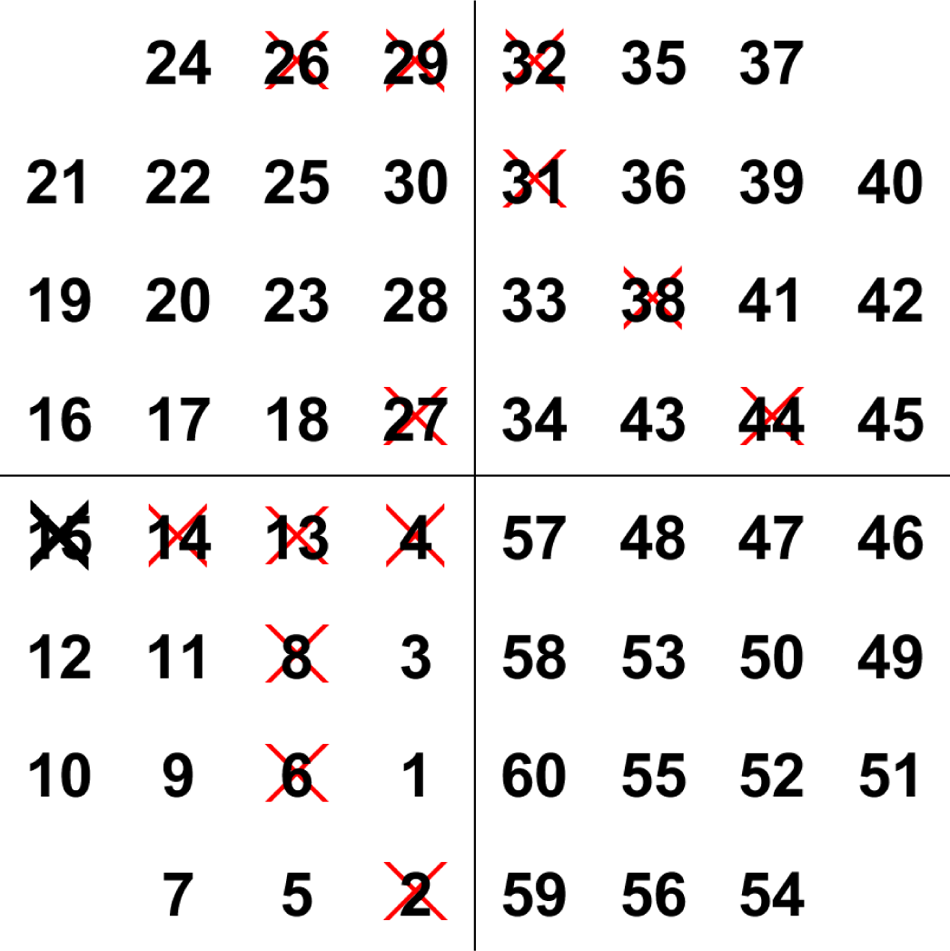
Schematic of the multielectrode array’s 59 electrodes, arranged in an 8-by-8 square grid with the 4 corners missing. Each electrode is identified by number. Being the ground electrode, electrode 15 is crossed out as it doesn’t detect electrical activity from the culture. This paper finds 13 defective electrodes, indicated by red crosses.

The full data as downloaded from the server maintained by Dr. Steve Potter’s laboratory at Georgia Institute of Technology is 86 gigabytes. We accepted their signal processing (including spike detection) performed using MeaBench [8]. We focus on *ℓ* = 920 recordings of daily spontaneous electrical activity from 58 cultures split across 8 batches, described in Table I. Each recording is associated with

**TABLE I.** Culture properties.

| Batches | Cultures | Plating density category | Age range (div) |
| --- | --- | --- | --- |
| 1 | 1-5 | dense | 5-25 |
| 2 | 1-6 | dense | 4-35 |
| 3 | 1-6 | dense | 7-31 |
| 4 | 1-2 | dense | 3-29 |
| 5 | 1-3 | dense | 5-39 |
|  | 4-7 | sparse | 5-40 |
|  | 8-10 | small | 5-39 |
| 6 | 1-3 | dense | 4-34 |
|  | 4-7 | sparse | 4-34 |
|  | 8-10 | small | 4-34 |
| 7 | 1-2 | dense | 4-35 |
|  | 3-6 | sparse | 4-35 |
|  | 7-10 | small | 4-35 |
| 8 | 1-3 | dense | 6-34 |
|  | 4-6 | small-and-sparse | 6-34 |
|  | 7-9 | ultra-sparse | 6-34 |

- a plating density category as defined in Table II,
- a batch (identifying a litter of embryos), numbered in experimental order,
- a culture (batch replicate plated on an MEA) number, and
- the culture developmental age (in *days in vitro*, div).

**TABLE II.** Plating density categorization definition. Five categories were defined based on measurements on a subset of cultures at 1 div[8].

| Category | Plating density (cells/mm<sup>2</sup>) |
| --- | --- |
| dense | 2500 ± 1500 |
| small | 1600 ± 600 |
| sparse | 600 ± 240 |
| small-and-sparse | 300 ± 160 |
| ultra-sparse | 110 ± 60 |

We analyzed these recordings using our own custom Matlab (The Mathworks, Natick, MA, USA) code. We treat each recording *i* as a sample and each electrode *j* as a feature describing the spike activity in each culture towards exploring spike patterns contained therein. The spike detection rate *SDR*_*i,j*_ is the number of detected spikes per second by electrode *j* during recording *i*. Following Wagenar’s lead [8], the array-wide spike detection rate

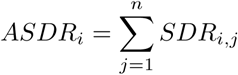

is the number of spikes detected across all electrodes per second.

The median ASDR is 55.7 spikes per second (interquartile range 12.9 − 158), but this distribution is strongly affected by many variables, e.g.:

- *Dense* cultures have a median ASDR of 118 spikes per second (IQR 49.6 − 233), or ≈ 17 times greater than the *ultra-sparse* median of 7.01 spikes per second (IQR 6.86 − 7.64).
- Older cultures (> 20 div) have a median ASDR of 141 spikes per second (IQR 48.6 − 257), or ≈ 12 times greater than younger cultures (< 10 div) median of 11.7 spikes per second (IQR 7.70 − 21.2).

These trends are consistent with those previously reported [8]. We consider

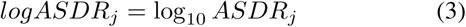

and *logASDR* to be a vector containing those values for all recordings, and

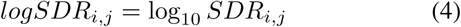

so that *logSDR*_*i,j*_ = log10 *SDR*_*i,j*_ (4) so that *logSDR*_*i,j*_ is the order of magnitude of the *j*^th^ electrode spike detection rate during the *i*^th^ recording. These values are assembled into matrix *logSDR* having *ℓ* rows (or recordings) and *n* columns (or electrodes). We also define

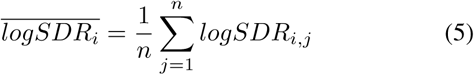

to be a recording’s mean *logSDR* across its electrodes and 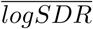to be a vector containing these averages for all recordings.

Since high correlations are observed across electrodes and across recordings, we perform dimensionality reduction on these data using principal components analysis (PCA) [18], [19]. PCA is an efficient linear transformation that creates new variables (called components) that are uncorrelated and sorted in decreasing order of variance. Each column of *logSDR* was mean-centered to generate matrix *X*, and the principal components (PCs) computed by means of SVD. The PC projections are the columns 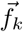 of *F* = *US*; these are plotted later in this paper (Figures 4-5). The relative contribution (expressed as a percentage) of PC *k* to the total variance is given by

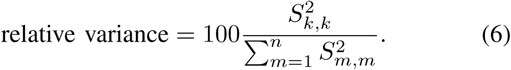

We examine the effect of the number of recording electrodes on our PCA representations by considering the cosine similarity between projections thereof. Consider the matrices *X*^[*n*]^ and *X*^[*m*]^. (Both matrices possess the same number of rows.) The similarity of both projections’ *k*^th^ components is given by

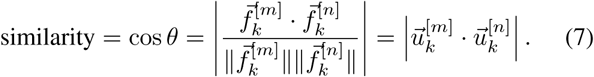

Angle *θ* is the angle between 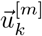 and 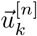. These similarities are plotted later in this paper, as Figure 6.

## IV. Results and Analysis

We present our results and their significance.

### A. Identifying systematic artifacts

In this first subsection, we establish that data contain two forms of defective electrodes that constitute systematic measurement error: **null electrodes** that failed to record any spikes over defined time intervals, and **weak electrodes** that sporadically under-recorded spike activity.

We begin by displaying matrix *logSDR* as the annotated heat map of Figure 2, with rows being recordings and columns being electrodes. Recordings are sorted by batch, then by culture, then by culture age. We observe the following features and trends:

**Fig. 2.**
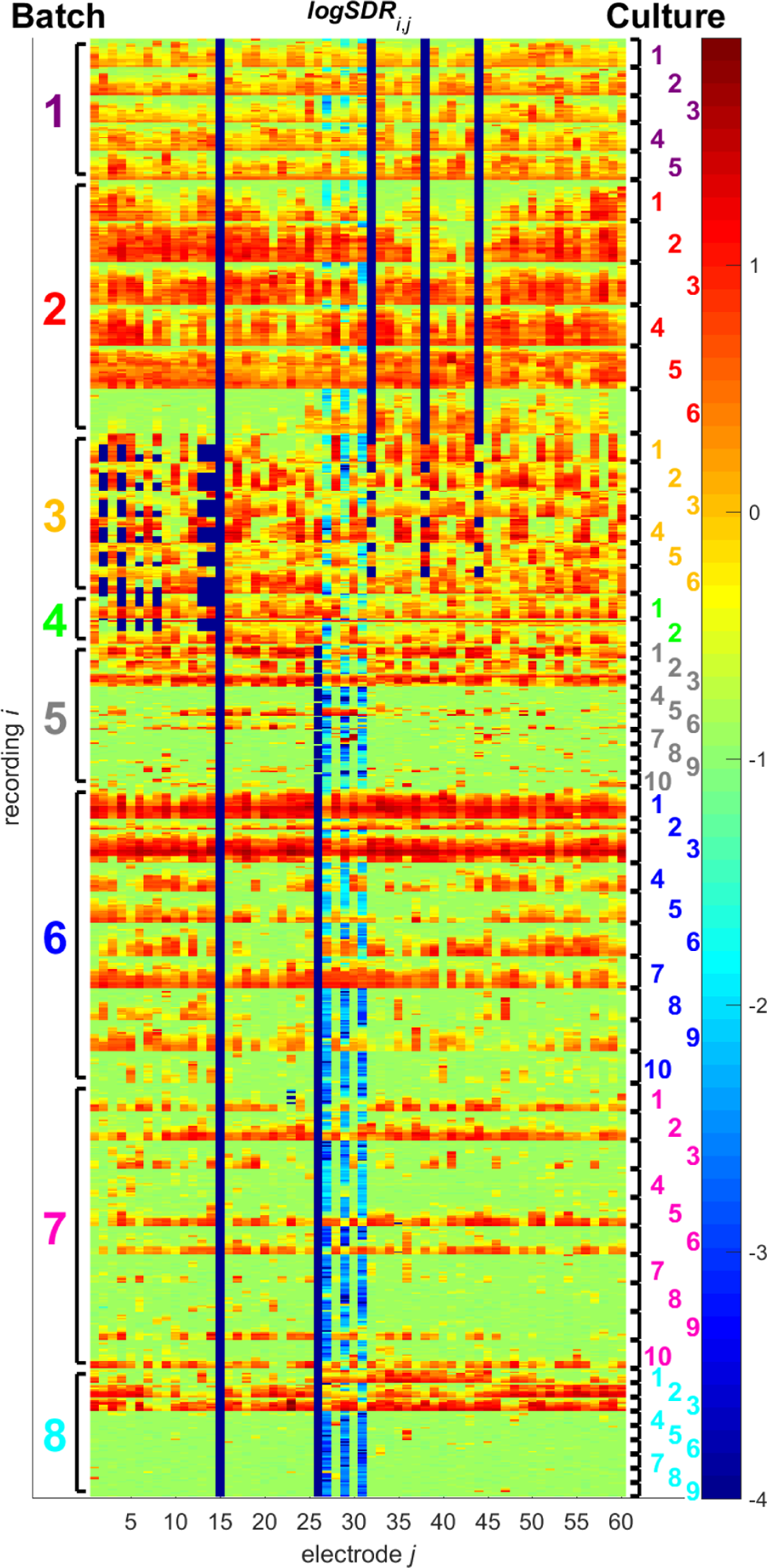
Heat map of the *logSDR* matrix displaying the order of magnitude of the spike detection rate (in spikes per second) at electrode *j* for recording *i*. The 920 recordings (rows) are sorted by batch number (identified on the left of the heat map), then by culture number (labeled on the right of the heat map), and finally by culture age. Entries of − ∞ (corresponding to zero spikes detected) were reset to − 4 so as to contrast with the next smallest value of − 3.07. This figure highlights ground electrode 15 (that didn’t participate in spike detection, full navy blue column), null electrodes 2, 4, 6, 8, 13, 14, 26, 32, 38, and 44 (that didn’t detect any spikes over specific time intervals, partial navy blue columns) and weak electrodes 27, 29, and 31 (that underdetected spikes throughout, lighter blue columns).

1. For a given recording, its *logSDR* is fairly homogenous across electrodes. Put otherwise, the inter-recording variability is greater than the inter-electrode variability.
2. Recordings from *dense* cultures frequently display *SDR* greater than 10 spikes/second (red). There is more red in the upper half of the heatmap (batches 1-4, all cultures plated dense). Cultures showing red in the lower half (batches 5-8, selected cultures plated dense) generally identify cultures plated at this highest density.
3. As expected, no spike was recorded from electrode 15, being the ground electrode. This appears in the heatmap as a navy blue column.
4. No spike was recorded for electrodes 32, 38, and 44 for all batch 1-2 recordings, and also for some batch 3 recordings. Similarly, no spike was recorded by electrode 26 for nearly all batch 5 recordings, and also for all batch 6-8 recordings. Patterns of spike non-detection are also seen in electrodes 2, 4, 6, 8, 13, and 14. These all appear as partial navy blue columns. We call these the **null electrodes**.
5. There appears to be an electrical activity *baseline* at ≈ 10^−1^ spike/second (green) that is seen in young cultures (whose cells were recently dissociated) and cultures plated at the lowest plating densities (batch 8 cultures 4-9). This baseline is more formally established in Figure 3.
6. In many recordings, electrodes 27, 29, and 31 appear to have under-detected spikes by at least 1 order of magnitude (light blue). These are the **weak electrodes**.

The presence of electrodes for which the recorded activity is significantly below the baseline of ≈ 10^−1^ spike/second (item 6) or for which no spike is recorded (item 4) are concerning. We fully document these null and weak electrode defectiveness artifacts in Table III.

**TABLE III.** Description of defective electrodes. This table documents the observed defective patterns in recording electrodes.

| Defective electrodes | Description of defectiveness | Affected recordings | Proportion of affected recordings (%) | Group explained |
| --- | --- | --- | --- | --- |
| 32, 38, 44 | Spike nondetection | Batches 1-2, and batch 3 to 17 div | 31.3 | A |
| 2 | Spike nondetection | Batch 3 from 18 div and batch 4 to 6 div | 7.0 | B |
| 4, 13, 14 | Spike nondetection | Batch 3 from 18 div and batch 4 to 15 div | 8.4 | B |
| 6, 8 | Spike nondetection | Batch 3 from 25 div and batch 4 to 15 div | 4.4 | B |
| 26 | Spike nondetection | Batch 5 from 6 div, and batches 6-8 | 57.3 | D |
| 27 | Spike underdetection | No clear pattern | 58.8 | n/a |
| 29 | Spike underdetection | No clear pattern | 57.0 | n/a |
| 31 | Spike underdetection | No clear pattern | 56.8 | n/a |
| 23 | Spike nondetection | Batch 7 culture 1 at 10, 14, 17, and 21 div | 0.4 | n/a |
| 35 | Spike nondetection | Batch 7 cultures 5-6 at 34 div | 0.2 | n/a |

**Fig. 3.**
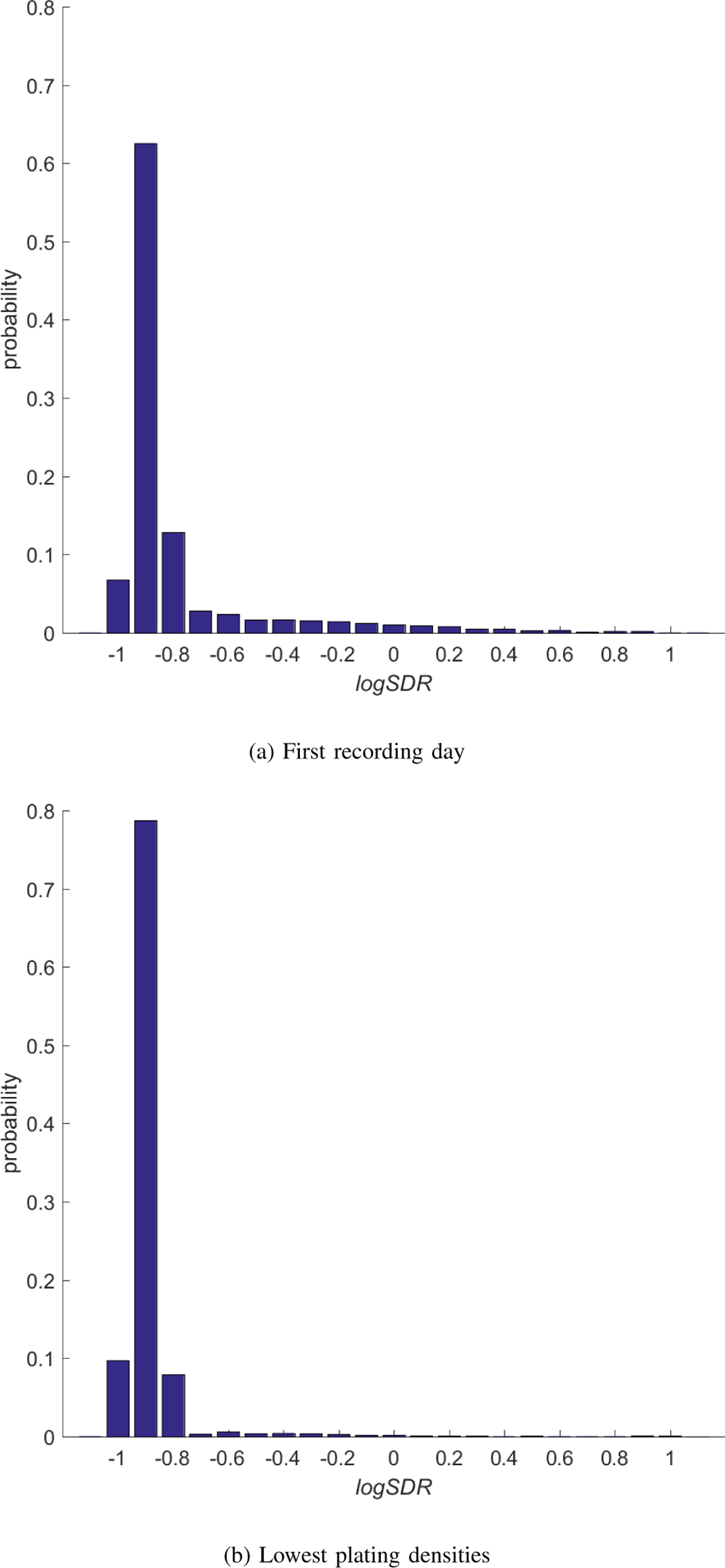
(a) Distribution of *logSDR* on the first recording day (identified in Table I) for all 58 cultures. Their cells having been recently dissociated, minimal electrical activity is seen. (b) Distribution of *logSDR* from all *small-and-sparse* and *ultra-sparse* recordings. These cultures didn’t show significant development, presumably due to an insufficient concentration of cells. Both distributions exclude null and weak electrodes. This figure establishes an electrical activity *baseline* of *logSDR* ≈ −1, or *SDR* ≈ 0.1 spike/second.

<u>The first portion of this table</u> defines four groups of null electrodes, each associated with a time interval:

- Group A: No spike was recorded for electrodes 32, 38, and 44 from the beginning of the experiments until 17 div of batch 3.
- Group B: No spike was recorded for electrodes 2, 4, 6, 8, 13, and 14 from 18 div of batch 3 until 15 div of batch 4. (There was some variation in start and end date for some of these electrodes, as fully documented in Table III.)
- Group C: All electrodes recorded normally starting on 16 div for batch 4 and lasting to 5 div of batch 5. (Since Table III documents defective electrodes, this group doesn’t explicitly appear in it.)
- Group D: No spike was recorded by electrode 26 from 6 div for batch 5 until the conclusion of the experiments.

Thereupon, through Figure 2 and Table III, we have identified and documented patterns for the following set of 10 null electrodes:

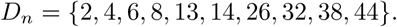

The proportion of recordings affected by each defective electrode appears in Table III.

<u>The second portion of Table III</u> documents 3 weak electrodes whose recorded level of signal activity is noticeably below that from other electrodes as seen in Figure 2 and mentioned in item 6. We consider a recording to have been affected by electrode 27, 29, or 31 if its *logSDR* is lesser than the smallest *logSDR* across the non-defective electrodes. All three electrodes are deemed to have under-detected spikes in a majority of recordings, compared to an expectation that such an electrode would have the smallest *logSDR* in 2.1% of recordings. We therefore declare the following set of 3 weak electrodes:

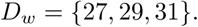

<u>The third portion of Table III</u> lists electrodes 23 and 35 as having not detected any spikes in four and two recordings, respectively. These defects do not appear to impact the placement of recordings in Figures 4-5, and are listed for completeness. In future analyses, these six *holes* in the *logSDR* matrix could be filled by interpolation, or their associated recordings thrown out.

**Fig. 4.**
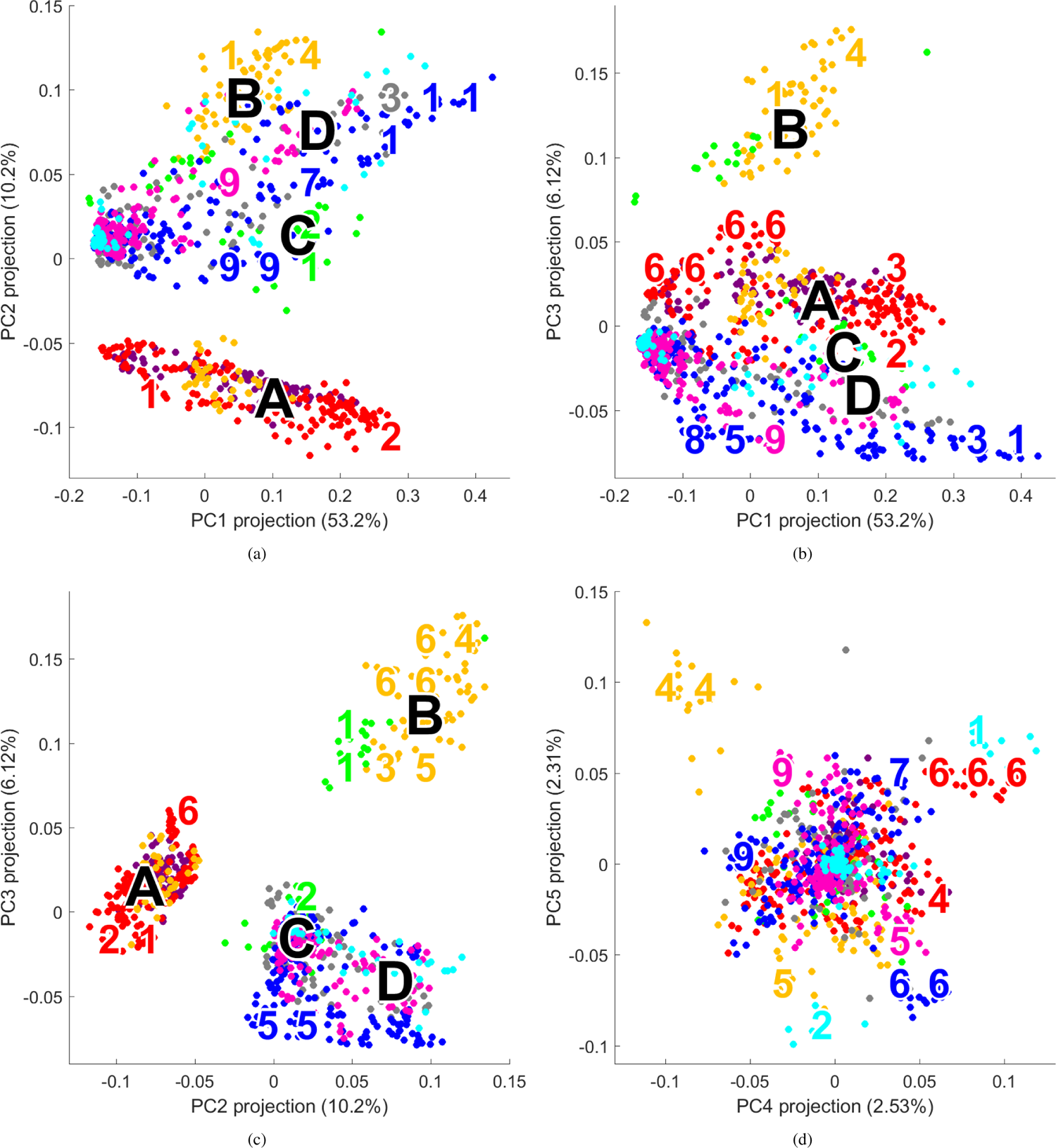
PCA biplot representations of spike detection rate across *n* = 59 electrodes. Panels a-c display three views of the leading 3 components. Panel d presents a different perspective: PC4 versus PC5. Each point is a recording. Colors denote batches: **Batch 1, Batch 2, Batch 3, Batch 4, Batch 5, Batch 6, Batch 7, Batch 8**. Regions associated with a specific culture are identified by culture number colored by batch. For instance, the the upper-left corner of panel d, we observe a concentration of plate 3 (gold) culture 4 recordings. As analyzed in the text: PC1 is highly correlated with logASDR; PC2 and PC3 discriminate 4 groups (labeled by letter) associated with defective electrodes; PC4 and beyond appear to distinguish specific cultures whose recordings present a nonuniform spike activity profile across electrodes.

Sets *D*_*n*_ and *D*_*w*_ document previously-unreported sources of systematic measurement error in these electrophysiological activity recordings. On average, each recording was affected by 1.9 null electrodes (standard deviation (SD) 1.3) and 1.7 weak electrodes (SD 1.2). Aggregating these, each recording was affected by an average of 3.7 defective electrodes (SD 1.6).

### B. Recognizing artifact impact on PCA

In this second subsection, we examine the PCA representations of the full *logSDR* matrix (Figure 4). The relative variance (equation 6) associated with each PC appears within axis labels. In doing so, we observe that defective electrodes introduce a heavy bias on these representations.

Typically, in PCA analyses, the PC1 projection is associated with overall (or average) sample magnitude, because this is the leading source of variance across the samples [20]. Indeed, our 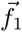 is almost perfectly correlated with 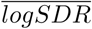 (Pearson’s correlation coefficient *r* = 0.998, *p* < 10^−6^). R<u>elating t</u>his finding back to array-wide metrics, we note that 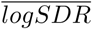 is highly correlated with *logASDR* (*r* = 0.960, *p* < 10^−6^). We therefore declare that the dominant trend (PC1, 53.2% of the variance) in matrix *logSDR* is the variation in overall spike activity across all recordings. Qualitatively, in Figures 4a and 4b, leftmost markers represent recordings showing minimal spike detection (in young cultures or those plated at the lowest densities) and rightmost markers represent highly-active recordings.

The second-greatest source of variance in matrix *logSDR* is the bias from null defectiveness patterns (PC2 and PC3, 16.3%). Indeed, Figure 4c discriminates Group A, B, and C/D associated with null electrode defectiveness:

- Group A includes all batch 1 (purple) and batch 2 (red) recordings, as well as some batch 3 (gold) recordings.
- Group B includes remaining batch 3 recordings and some batch 4 (lime) recordings.
- Group C includes remaining batch 4 recordings.
- Group D includes all batch 5 (grey), batch 6 (blue), batch 7 (pink) and batch 8 (green) recordings.

These groups are so labeled in Figure 4. Groups A and B separate better because they are each defined by several defective electrodes. Groups C and D are intermixed as they differ by a single null electrode (electrode 26, as previously documented).

The third-greatest source of variance identifies characteristics proper to individual cultures in PC4 and higher (represented as colored numbers appearing on the edges of the cloud in Figure 4d). Most visibly, all recordings from batch 3 (gold) culture 4 appear in the upper-left region of Figure 4d.

### C. Generating unbiased PCA model

In this third subsection, we remove the 13 columns of *logSDR* associated with the defective electrodes in set *D*_*n*_ ∪ *D*_*w*_, so that the number of electrodes is now *n* = 46. In doing so, we expect that the 4 groups appearing in Figure 4 will disappear. Indeed, they do, and the regenerated PCA biplots appear as Figure 5.

**Fig. 5.**
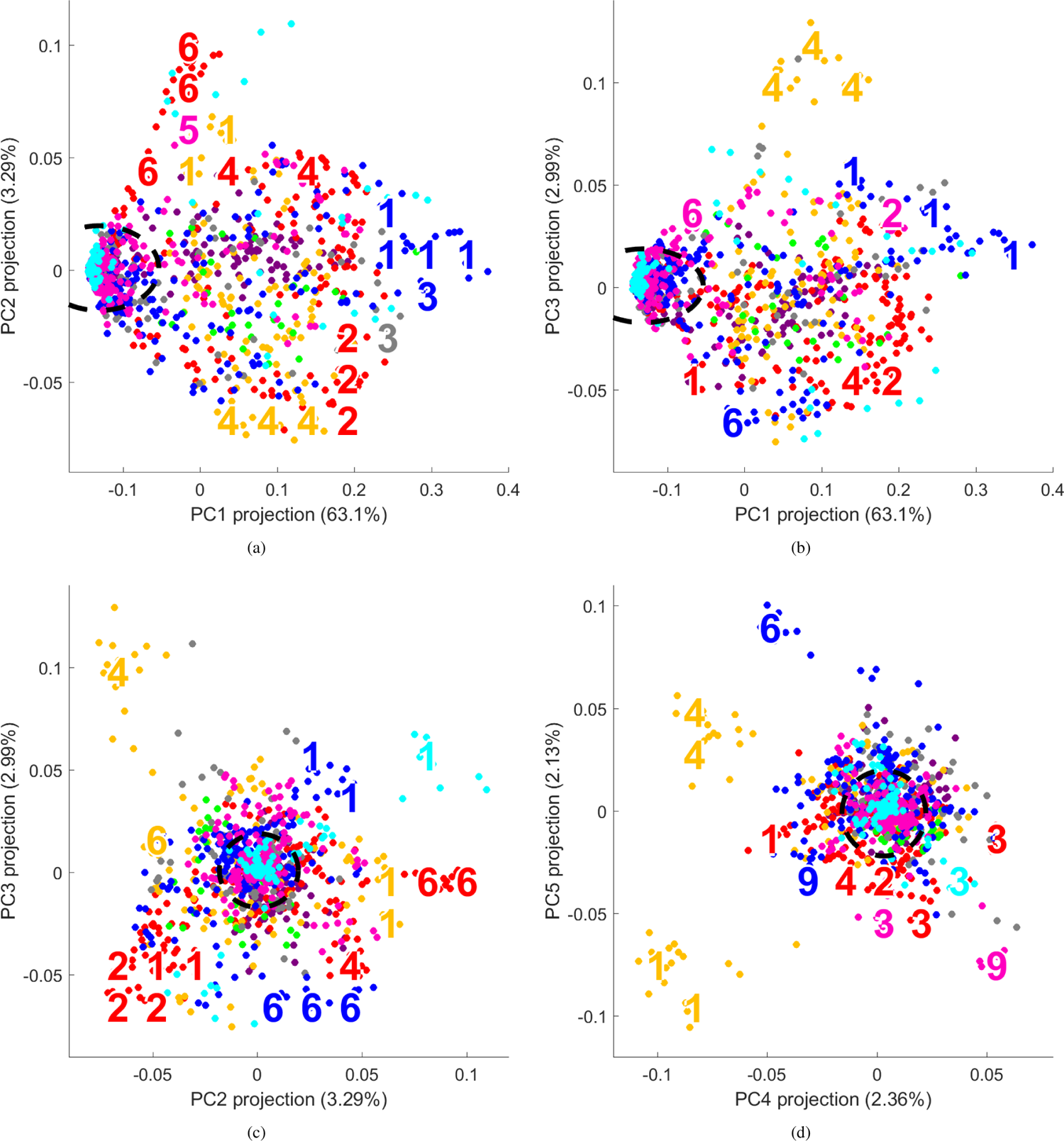
PCA biplot representations of spike detection rate across *n* = 46 electrodes, after removal of 13 defective electrodes. The ellipse denotes a region associated with minimally-active recordings. Other features of these biplots are consistent with the caption to Figure 4. As analyzed in the text: PC1 continues to be highly correlated with logASDR; the 4 defectiveness groups are no longer identified; PC2 and beyond appear to distinguish specific cultures (most visibly, batch 3 (gold) cultures 4 and 1).

The dominant source of variance in matrix *logSDR* remains spike detection magnitude, with PC1 being highly correlated with *logASDR* (Pearson’s correlation coefficient *r* > 0.999, *p* < 10^−6^) than before. Thus, the array-wide spike activity magnitude explains 63.1% of the dataset’s variability, with differences across individual electrodes accounting for the remainder. Inspection of the rightmost portion of Figures 5a and 5b identifies some of the most-active cultures: batch 5 (blue) cultures 1 and 3, and batch 5 (grey) culture 3.

By virtue of its interpretation, PC1 filters out the level of spike activity in each recording, so that the other components reflect patterns in spike activity across the electrodes that are scale-independent. Put otherwise, Figure 5c and 5d say nothing about *how many* spikes were detected in the recordings, but rather express similarities and differences in recordings’ spike distribution across the electrodes.

At 3.29%, PC2 accounts for less than one-tenth of the remaining variance. Indeed, several low-variance components (each within the 2-4% range) seem to distinguish specific individual cultures, starting with PC2. In these biplots, a number indicates a concentration of recordings belonging to that culture whose batch is identified by color. Some of the most strikingly identifiable cultures are:

- All recordings from batch 3 (gold) culture 4 appear in the upper-left region Figure 4c, and then again in the upper-right region of Figure 4d.
- All recordings from batch 8 (green) culture 1 appear in the upper-right region of Figure 4c.
- All recordings from batch 3 (gold) culture 1 appear in the lower-right region of Figure 4d.
- All recordings from batch 6 (blue) culture 6 appear in the lower-mid region of Figure 4c, and then again in the upper-mid region of Figure 4d.

These cultures each present a unique profile of baseline and highly-active spike detections across their electrodes in the Figure 2 heatmap.

Qualitatively, Figures 4d and 5c are strikingly similar, with several unique cultures positioned in the same locations (including batch 3 (gold) culture 4, batch 8 (green) culture 1, batch 2 (red) culture 6, and batch 6 (blue) culture 6). Removing the defective electrodes from our data eliminates the two components expressing that bias. In mathematical terms, the bias patterns are fairly orthogonal (perpendicular) to the unbiased trends found in these data.

### D. Effect of eliminated electrodes

Since we reduced our dataset from 59 to 46 electrodes, we explore the loss associated with those 13 defective electrodes on our analysis. We do this by computing the cosine similarity (equation 7) of a PC projection determined from *X*^[46]^ with the PC projection obtained from a reduced *X*^[*m*]^ consisting of an *m*-column subset of *X*^[46]^ chosen at random. By repeating this procedure 100 times and plotting the resulting distributions in Figure 6, we can quantify this loss.

**Fig. 6.**
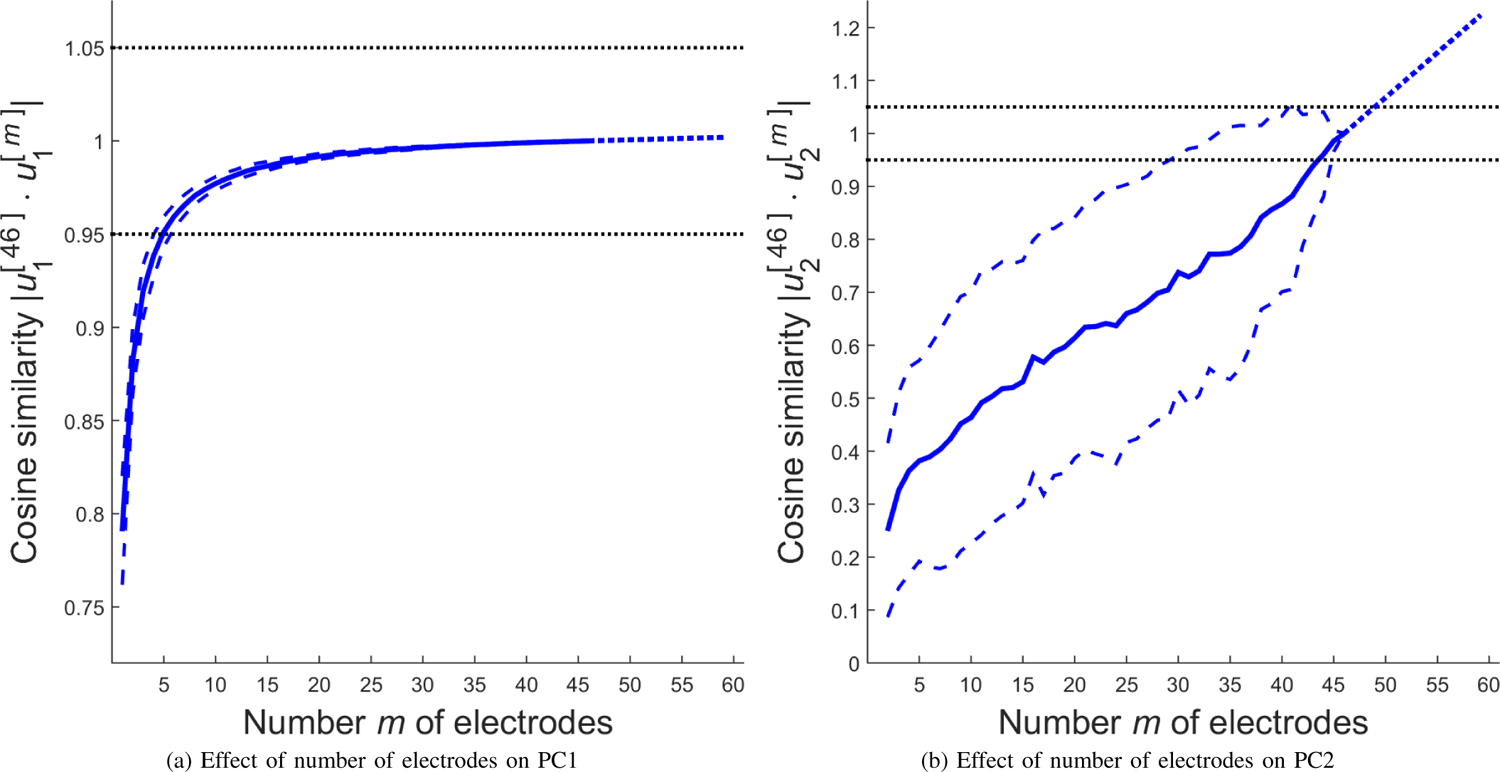
Effect of electrode count *m* on leading PC components. The mean cosine similarity (equation 7) to the 46-electrode model is plotted (solid curve). This curve is linearly extrapolated (dotted) to 59 electrodes. One standard deviation from the mean is also indicated (dashed curves). The 5% dissimilarity interval about the 46-electrode model also appears. (a) PC1 is robust to varying electrode count. A small number of electrodes is sufficient to capture PC1’s trend. Losing 13 electrodes had a minimal impact on its definition. (b) PC2 is highly sensitive to varying electrode count. Losing 13 electrodes has a great impact on its definition. Further, even 59 electrodes may have been insufficient to stabilize a definition for PC2.

We find that PC1 is robust to varying electrode count, whereas PC2 and higher components are sensitive to the number of electrodes involved.

Indeed, as few as 5 electrodes are sufficient to expect PC1 projections that are at most 5% dissimilar from the 46-electrode model. Linearly extrapolating (dotted line) to 59 electrodes predicts PC1 projections that are < 0.2% dissimilar from the 46-electrode model. (This extrapolation is probably pessimistic as the curve appears to converge.) The loss of 13 electrodes had a minimal impact on PC1. This is consistent with our interpretation that PC1 captures array-wide spike detection magnitude: only a few electrodes are sufficient to determine this level of activity.

On the flipside, adding or removing 3 electrodes is sufficient to expect PC2 projections that are more than 5% dissimilar from the 46-electrode model. Linearly extrapolating to 59 electrodes predicts PC2 projections that are 22% dissimilar from the 46-electrode model. The loss of 13 electrodes has a significant impact on our ability to interpret PC2 with confidence. Moreover, the lack of curvature in the trend also prevents us from speculating as to how many recording electrodes would be needed to stabilize a definition for PC2. In the context that PC2 participates in distinguishing specific cultures, it seems that those patterns are weak enough that they are less apparent with a few missing electrodes. We produced similar graphs for subsequent PCs (not shown) that are similar to Figure 6.

## V. Discussion

We begin this discussion by recalling that PC1 explains 63.1% (or roughly 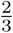 of the variance (Figure 5) in our *logSDR* matrix, and this component equates almost exactly to the *logASDR* variable. This first component that corresponds to array-wide spike activity magnitude, is the first *modality* of these data. This component is also robust, being accurately determined from a small number of electrodes as seen in our Figure 6a analysis. This finding supports a current development of high-throughput platforms utilizing multiple cell culture chambers with only a few electrodes per chamber. As long as *ASDR* is the primary metric for characterizing cultures, a few electrodes are sufficient, and our findings support the use of these multiwell designs. But, if we come to discover that richer metrics reveal additional meaningful insights, then more electrodes and/or alternate electrode arrangements may be needed.

Do this paper’s insights negate Wagenaar et al.’s [8] findings? Probably not, as their results were based on spike quantity. Indeed, *ASDR* would be mildly sensitive to the number of electrodes contributing to the sum; indeed, the number of defective electrodes differs by 1.7 on average (or 2.8% of 59 electrodes) across all recordings. Figure 7a exemplifies how this paper maintains previous findings based upon array-wide measures. Indeed, this figure swaps peak *log*_10_*ASDR* for peak PC1 projection and is almost identical to Figure 5b from Wagenaar et al. [8], reinforcing the fact that fewer spikes are recorded in lower-density cell cultures.

**Fig. 7.**
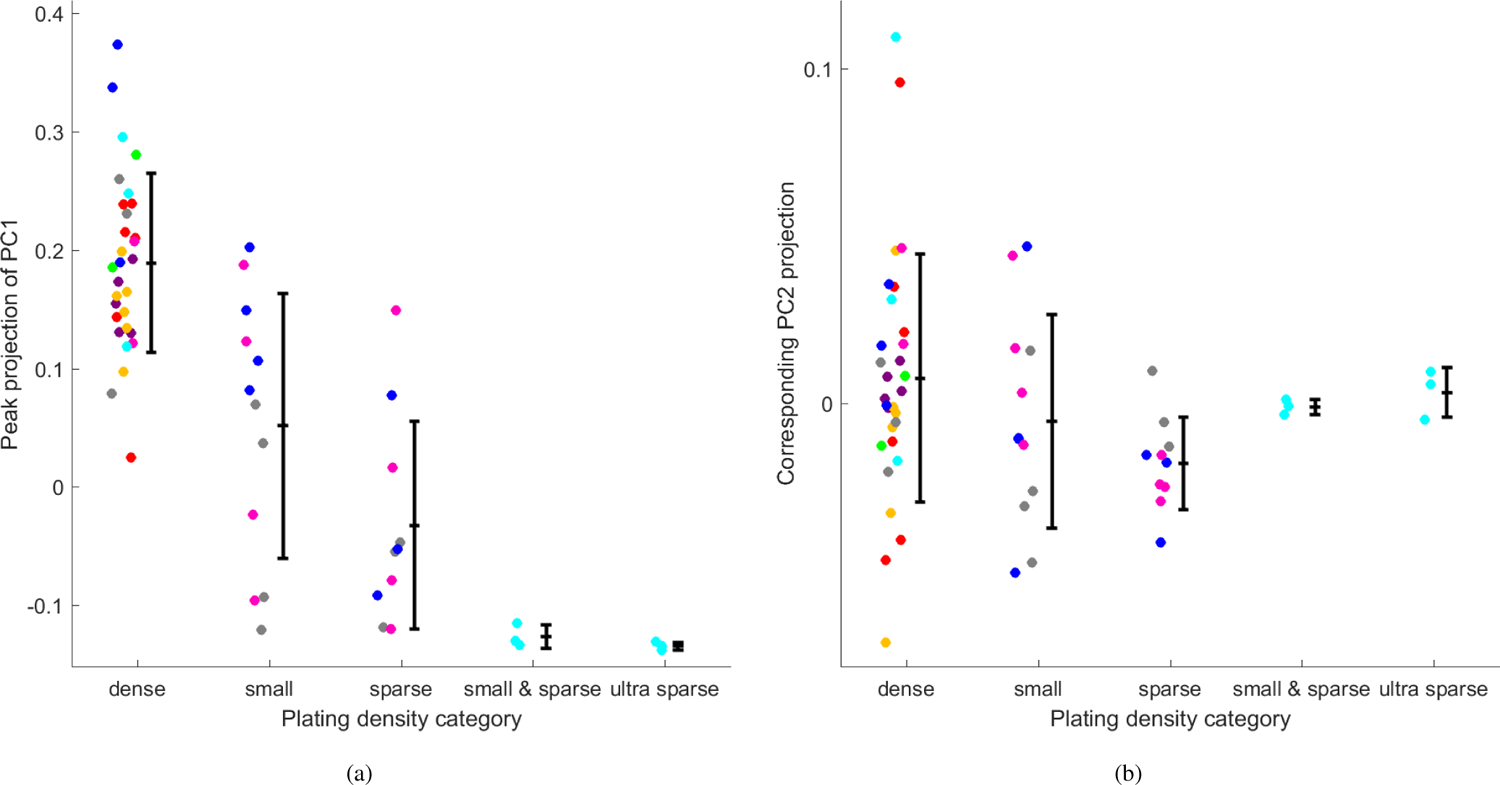
Each point represents a culture, on the recording day associated with its greatest PC1 projection. Error bars indicate the mean and standard deviation across all cultures plated within a density category. Dots were horizontally jittered for visual clarity. (a) Spike activity (or PC1) decreases with decreasing plating density. This replicates Figure 5b in Wagenaar et al. [8] that plotted log_10_ *ASDR* against the five plating density categories. This supports our view that this paper doesn’t negate array-wide trends reported in [8]. (b) PC2 is not strongly associated with plating density.

Further, several Wagenaar et al.’s figures [8] focus on specific batches or plating densities. We recall that most batches (all except 3-4) were each generally contained within a defective electrode grouping (Table III). We also recall that variations in plating density category were limited to batch 5-8 cultures that all belong to Group D. Thus, this paper’s findings don’t impact most of their intra-batch comparisons, nor all of their extra-category comparisons. Other studies shouldn’t be significantly impacted, e.g. [14], [15] focus solely on batch 2 cultures which is wholly contained within Group A.

In this paper, the bias introduced by defective electrodes is greater, because of our focus on *which electrodes* contributed to a metric, as is the case with our PCA representations (Figure 4), as opposed to *how many electrodes* contribute to a metric. Some other analyses of these data [12], [13], [16] may need to be revisited in light of our findings. Also, an awareness of the issues highlighted in this paper may apply to the analysis of other MEA datasets, of which only a few are listed here: [1], [2], [4]–[7].

By treating each electrode as a variable *SDR*_*j*_, rather than aggregating all electrodes into a single variable *ASDR*, we access the remaining 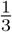 of the variance that hopefully reveals insights into cortical culture spike activity. Most recordings’ spikes are distributed uniformly across the electrodes, and these appear near the origin (black dashed ellipses) in Figures 5c and 5d. A few cultures possess recordings whose spike distributions are unique; these cultures find themselves on the edges of Figure 5c or 5d. Related to this, our analysis of Figure 6b found that these unique patterns are sensitive to a few electrodes being added or removed, suggesting that these differences in development patterns are subtle. Principal components 2-5 also each account for a meager proportion of the overall variability. We had hoped that PC2 and subsequent components might reveal richer *modalities* that would transcend batches, plating densities and culture age. Figure 7b exemplifies this lack of association between PC2 and plating density. Those modalities may still exist, but may not be revealed until the PC definitions stabilize with more electrodes. At the very least, our underwhelming interpretation for PC2-PC5 speaks to the challenges of extracting higher-order knowledge from MEA data, and perhaps validates the widespread use of *ASDR*.

Yet, a few reasons lead us to believe that a deeper analysis of these data could yield new insights.

1. Figure 5 presents a curious phenomenon, with intra-batch variability dominating over inter-batch variability. Most visibly, recordings corresponding to batch 2 (red) cultures (that were all plated *dense*) are scattered across the biplots, with no indication that those recordings are more similar to each other than to recordings from other batches (or plating densities).
2. Bartlett’s test [21] suggests that 40 principal components are necessary to explain the nonrandom variation in *logSDR* at the *α* = 0.05 significance level. Implementing supervised learning techniques such as support vector machines (SVM) [22] following a robust experimental design [23] may therefore be required to uncover those insights.
3. Sought patterns may not appear in linear space. The SVM could identify these nonlinear features, but their visualization would require kernel PCA [24] or independent components analysis [25].
4. Ultimately, we are interested in transitioning from quantity metrics (such as SDR) towards measuring spatial relationships for describing tissue interconnectivity from MEA data. This is a challenging problem, as our preliminary efforts to do this showed limited enrichment over SDR [26].

It’s important to note that the physical locations of the defective electrodes, per Figure 1, bear some relation to the groups defined in Table III. Further, these groups roughly correspond to physical features of the recording system itself. Cables connecting the preamplifier to the recording computer divide the recorded electrodes along lines roughly following the quadrants of Figure 1. As it turns out, all Group B electrodes are associated with a particular cable and breakout box. Similarly, all weak electrodes are associated with a different cable and breakout box. Furthermore, two of the three Group A electrodes share a third distinct physical connection. This brings another caution to the foreground. While these aspects are unique to Potter’s system, which may be subject to more unit-to-unit variation, similar factors can have an impact on the reliability of such recordings. In a follow-up experiment, Wagenaar et al. glean an awareness of the issues raised in this paper, and offer an explanation for why an electrode’s spikes might not be recorded: *due to a broken wire in one pre-amplifier channel, only 58 [electrodes] could be used for recording* [11].

Naïve observations of Figure 4 would have lead to the premature conclusion that these recordings provide sufficient information to distinguish between specific batches. This fits nicely with our objectives of identifying inherent biological attributes that are retained in vitro. Had plate 3 (gold) not been split across Groups A and B, and had plate 4 (lime) not been split across Groups B and C, we might have not realized so quickly that these groups were a systematic measurement error manifestation.

This situation motivates the importance of best practice protocols for analyzing MEA data, that we propose as follows:

1. Possessing a system of procedures and standards for identifying and documenting artifacts in MEA data, so that shared data is annotated in a meaningful manner.
2. Regularly testing all electrode connections.
3. Visualizing recordings sorted chronologically (similar to Figure 2) to seek null and weak electrode patterns.
4. Create PCA representations (similar to Figure 4) towards identifying more subtle patterns of electrode defectiveness.

This review process is critical prior to applying data analysis algorithms [27]–[31]. However, we go beyond the identification of two forms of electrode defectiveness; we also demonstrate possibilities for post-analytic review. Indeed, removing the affected electrodes eliminated the bias caused by measurement error, as seen by comparing Figures 4-5.

## VI. Conclusion

To quote the first sentence of Kello et al.: *Complexity is widespread in neuronal spike trains and propagation of spike activity* [31]. We presented a protocol towards ensuring that this complexity is not further muddled by systematic measurement error. We also demonstrated how proper data curation is critical towards gaining data analytic results that are meaningful.

We demonstrated possibilities for post-experiment data analytic review towards identifying new and unknown issues while recording, and retroactively handle those issues. From here, we aim to better identify and interpret the associations and correlations found in Figure 5, including those in higher-order components. Ultimately, biased variability as a result of defective electrodes can be avoided wholly by proper documentation of experimental conditions. This highlights the need for best practices when recording MEA signals.

Our next steps involve digging deeper into the interpretation of our principal components beyond PC1, and thinking about how to transition from a paradigm where each recording is a sample to one where each culture is a sample, thereby permitting us to characterize cell culture development.

## Acknowledgment

The authors thank Daniel Wagenaar, Jerome Pine and Steve Potter for making their data available and encourage their post-analytic review [8]. We also thank the Albany College of Pharmacy and Health Sciences *Scholarship of Discovery* program for funding a portion of this work.

## References

[1] A. J. Cadotte, T. B. DeMarse, P. He, and M. Ding, “Causal measures of structure and plasticity in simulated and living neural networks,” PLoS ONE, vol. 3, no. 10, p. e3355, 2008. [Online]. Available: http://dx.doi.org/10.1371%2Fjournal.pone.0003355

[2] S. Illes, S. Theiss, H.-P. Hartung, M. Siebler, and M. Dihné, “Niche-dependent development of functional neuronal networks from embryonic stem cell-derived neural populations,” BMC Neuroscience, vol. 10, no. 1, p. 93, 2009. [Online]. Available: http://www.biomedcentral.com/1471-2202/10/93

[3] A. F. Johnstone, G. W. Gross, D. G. Weiss, O. H.-U. Schroeder Gramowski, and T. J. Shafer, “Microelectrode arrays: A physiologically based neurotoxicity testing platform for the 21st century,” NeuroToxicology, vol. 31, no. 4, pp. 331–350, 2010. [Online]. Available: http://www.sciencedirect.com/science/article/pii/S0161813X10000677

[4] E. Biffi, G. Regalia, A. Menegon, G. Ferrigno, and A. Pedrocchi, “The influence of neuronal density and maturation on network activity of hippocampal cell cultures: A methodological study,” PLoS ONE, vol. 8, no. 12, p. e83899, 12 2013. [Online]. Available: http://dx.doi.org/10.1371%2Fjournal.pone.0083899

[5] P. Massobrio, C. Giachello, M. Ghirardi, and S. Martinoia, “Selective modulation of chemical and electrical synapses of helix neuronal networks during in vitro development,” BMC Neuroscience, vol. 14, no. 1, p. 22, 2013. [Online]. Available: http://www.biomedcentral.com/1471-2202/14/22

[6] M. Bisio, A. Bosca, V. Pasquale, L. Berdondini, and M. Chiappalone, “Emergence of bursting activity in connected neuronal sub-populations,” PLoS ONE, vol. 9, no. 9, p. e107400, 09 2014. [Online]. Available: http://dx.doi.org/10.1371%2Fjournal.pone.0107400

[7] M. Hammond, D. Xydas, J. Downes, G. Bucci, V. Becerra, K. Warwick, A. Constanti, S. Nasuto, and B. Whalley, “Endogenous cholinergic tone modulates spontaneous network level neuronal activity in primary cortical cultures grown on multi-electrode arrays,” BMC Neuroscience, vol. 14, no. 1, p. 38, 2013. [Online]. Available: http://www.biomedcentral.com/1471-2202/14/38

[8] D. Wagenaar, J. Pine, and S. Potter, “An extremely rich repertoire of bursting patterns during the development of cortical cultures,” BMC Neuroscience, vol. 7, no. 1, p. 11, 2006. [Online]. Available: http://www.biomedcentral.com/1471-2202/7/11

[9] D. A. Wagenaar, R. Madhavan, J. Pine, and S. M. Potter, “Controlling bursting in cortical cultures with closed-loop multi-electrode stimulation,” J Neurosci, vol. 25, no. 3, pp. 680–688, 2005.

[10] D. Wagenaar, T. DeMarse, and S. Potter, “Meabench: A toolset for multielectrode data acquisition and on-line analysis,” in Neural Engineering, 2005. Conference Proceedings. 2nd International IEEE EMBS Conference on, March 2005, pp. 518–521.

[11] D. Wagenaar, J. Pine, and S. Potter, “Searching for plasticity in dissociated cortical cultures on multi-electrode arrays,” Journal of Negative Results in BioMedicine, vol. 5, no. 1, p. 16, 2006. [Online]. Available: http://www.jnrbm.com/content/5/1/16

[12] D. A. Wagenaar, Z. Nadasdy, and S. M. Potter, “Persistent dynamic attractors in activity patterns of cultured neuronal networks.” Phys Rev E Stat Nonlin Soft Matter Phys, vol. 73, no. 5 Pt 1, May 2006. [Online]. Available: http://view.ncbi.nlm.nih.gov/pubmed/16802967

[13] J. D. Rolston, D. A. Wagenaar, and S. M. Potter, “Precisely timed spatiotemporal patterns of neural activity in dissociated cortical cultures,” Neuroscience, vol. 1, no. 148, pp. 294–303, Aug. 2007. [Online]. Available: http://www.ncbi.nlm.nih.gov/pubmed/17614210/

[14] D. Patnaik, P. Sastry, and K. Unnikrishnan, “Inferring neuronal network connectivity from spike data: A temporal data mining approach,” Journal of Scientific Programming, vol. 16, pp. 49–77, 2008.

[15] D. Patnaik, S. Laxman, and N. Ramakrishnan, “Discovering excitatory relationships using dynamic bayesian networks,” Knowledge and Information Systems, pp. 1–31, 2010, 10.1007/s10115-010-0344-6. [Online]. Available: http://dx.doi.org/10.1007/s10115-010-0344-6

[16] A. Napoli, J. Xie, and I. Obeid, “Understanding the temporal evolution of neuronal connectivity in cultured networks using statistical analysis,” BMC Neuroscience, vol. 15, no. 1, p. 17, 2014. [Online]. Available: http://www.biomedcentral.com/1471-2202/15/17

[17] G. H. Golub and C. F. Van Loan, Matrix Computations, 3rd ed. Baltimore: Johns Hopkins University Press, 1996.

[18] I. T. Jolliffe, Principal Component Analysis, 2nd ed. New York: Springer-Verlag, 2002.

[19] H. Abdi and L. J. Williams, “Principal component analysis,” Wiley Interdisciplinary Reviews: Computational Statistics, vol. 2, no. 4, pp. 433–459, 2010. [Online]. Available: http://dx.doi.org/10.1002/wics.101

[20] J. O. Ramsay and B. W. Silverman, Functional Data Analysis, 2nd ed. New York: Springer, 2005.

[21] J. H. McDonald, Handbook of Biological Statistics, 3rd ed. Baltimore: Sparky House, 2014.

[22] N. Cristianini and J. Shawe-Taylor, Support vector machines and other kernel-based learning methods. Cambridge University Press, 2000.

[23] S. Arlot, “A survey of cross-validation procedures for model selection,” Statistics Surveys, vol. 4, pp. 40–79, 2010.

[24] J. Shawe-Taylor and N. Cristianini, Kernel Methods for Pattern Analysis. Cambridge, UK: Cambridge University Press, 2004.

[25] A. Hyvärinen, J. Karhunen, and E. Oja, Independent Component Analysis, ser. Adaptive and Cognitive Dynamic Systems: Signal Processing, Learning, Communications and Control. Wiley, 2004. [Online]. Available: http://books.google.com/books?id=96D0ypDwAkkC

[26] T. Stone, U. Khawaja, N. Perko, T. Kiehl, and C. Bergeron, “Best practices for avoiding dominant experimental bias in analysis of multielectrode array signals,” BMC Neuroscience, vol. 15, no. Suppl 1, p. P208, 2014. [Online]. Available: http://www.biomedcentral.com/1471-2202/15/S1/P208

[27] A. Barrat, M. Barthelemy, R. Pastor-Satorras, and A. Vespignani, “The architecture of complex weighted networks,” P Natl Acad Sci USA, vol. 101, no. 11, pp. 3747–3752, 2004.

[28] E. Brown, R. Kass, and P. Mitra, “Multiple neural spike train data analysis: state-of-the-art and future challenges,” Nat Neurosci, vol. 7, pp. 456–461, 2004.

[29] E. Bullmore and O. Sporns, “Complex brain networks: graph theoretical analysis of structural and functional systems,” Nat Rev Neurosci, vol. 10, pp. 186–198, 2009.

[30] G. Fogel, “Computational intelligence approaches for pattern discovery in biological systems,” Briefings in Bioinformatics, vol. 9, pp. 307–316, 2008.

[31] C. T. Kello, J. Rodny, A. S. Warlaumont, and D. C. Noelle, “Plasticity, learning, and complexity in spiking networks,” Crit Rev Biomed Eng, vol. 40, no. 6, pp. 501–518, 2012.

